# Falcons use wind assistance and remote islands to mitigate risk during ocean-crossings

**DOI:** 10.1101/2025.08.12.669758

**Authors:** Meixu Chen, Duarte S. Viana, Jordi Figuerola, Laura Gangoso, Wouter M.G. Vansteelant

## Abstract

Wind conditions play a major role in determining how birds negotiate ecological barriers during long-distance migration. Migrants typically minimize flights over barriers in opposing winds, while crossing the same barriers directly in supportive winds. Nevertheless, even in supportive conditions, barrier-crossings likely involve a number of risk-mitigating behaviors, including the use of stepping stones. We aimed to understand how seasonal ocean-crossings of Eleonora’s falcons (*Falco eleonorae*) compare in terms of wind support and flight effort, and whether falcons use islands to cross safely in variable wind conditions. To do this we combined GPS-tracking data from 19 individuals over a decade (2012-2022) with wind data from an atmospheric reanalysis model. Despite major differences in wind support, falcons achieved similar ground speeds in both seasons. That is because falcons reduce instantaneous flight effort by flying at lower airspeeds in supportive spring winds (31.5 ± 6.0 km h^-1^) than in adverse autumn winds (47.4 ± 9.3 km h^-1^). Overall, it took falcons twice as many flight hours to complete the spring crossings that were 40% greater in terms of air distance compared to autumn crossings. Islands were more frequently used during spring crossings (57.9% individuals in spring vs 21.1% in autumn), and the probability of a falcon using an island significantly increased with weaker wind support in spring (but not in autumn). While wind support partially offsets the extra distance flown over the ocean in spring, isolated islands offer emergency stop-over opportunities when Eleonora’s falcons experience relatively weak wind support during protracted ocean crossings.

## Introduction

Long-distance migration often requires individuals to overcome ecological barriers; i.e. landscapes far outside a species’ habitual niche that offer limited opportunities for resting and refueling, or characterized by extreme environmental conditions (Alerstam, 1993; Lathouwers et al. 2022; Newton, 2023). For migrant landbirds, the riskiest barriers are large water bodies where they cannot land and thus face a risk of drowning in case of inclement weather or fatigue. As such, birds often circumvent large water bodies entirely, or minimize the time and distance flown over water by crossing water bodies at their narrowest points (e.g. sea-straits). The aversion for crossing water bodies is especially pronounced in obligate-soaring migrants (Mellone et al.,2020) that face disproportionately high flight costs when atmospheric conditions force them to resort to active, flapping flight - which is often the case over open water (but see: Duriez et al., 2018; Nourani et al., 2020, 2021; Pekarsky et al., 2024). More broadly speaking, the extent to which landbirds avoid water bodies depends in large part on prevailing winds: birds are more likely to cross directly in supportive than adverse winds (Gill et al 2014; Becciu et al 2020; Norevik et al 2020; Ciković et al., 2021; Bathrick et al 2024).

Wind support has such a strong effect on flight costs(Liechti, 2006) that prevailing winds explain the seasonal configuration of global aerial flyways in general (Kranstauber et al., 2015), and especially in relation to ecological barriers(Mellone et al., 2016; Newton, 2023; Shamoun-Baranes et al 2017). Indeed, numerous migrants exhibit seasonal loop migrations because prevailing winds (or other weather factors) permit them to cross a barrier directly in one but not the other season(Becciu et al., 2020; Nourani et al., 2016, 2021; Vansteelant, Kekkonen, et al., 2017; Vansteelant et al., 2021). Nevertheless, even with supportive winds, crossing a large expanse of open water in sustained active flight is not without risk. And so barrier-crossings likely involve a range of risk-mitigating behaviours, such as airspeed and flight altitude adjustments that can help reduce energy expenditure and increase a bird’s flight time and range (Blas et al., 2020; Literák et al., 2021; Vansteelant, Shamoun-Baranes, McLaren, et al., 2017; Norevik et al. 2023).

Another behavior by which birds could mitigate the risk of barrier-crossings is the use of isolated patches of more benign habitat. For example, recent tracking studies have shown that some Afro-Palearctic migrants make multi-day stop-overs while crossing the “sea of sand” of the Sahara Desert, suggesting that (in some years) certain parts of the desert actually do offer rest and foraging opportunities (Monti et al., 2024; Strandberg et al., 2010). Shorebirds crossing the “sea of green” of the Amazon Basin were found to use river banks and lakes in unexpectedly high numbers (Linscott et al., 2024), while forest patches serve as stepping stones for landbirds traversing the Corn Belt in the United States (Guo et al., 2025). Analogously, it seems only logical that ocean-crossing landbirds would employ an island-hopping strategy, as they appear to do in the Central Mediterranean and along island chains in eastern Asia (Agostini et al., 2004; Bildstein, 2006; Nourani et al., 2018). Yet, it is unclear what role remote islands have in moulding trans-oceanic migrations of landbirds more generally, and if so, whether they are used on a regular basis as stepping stones, or more opportunistically as “emergency stop-overs” (Shamoun-Baranes et al., 2010). Inclement weather during sea-crossings can result in mass mortality events (Newton, 2024) and so-called fall-outs whereby huge numbers of birds land on islands, offshore structures and ships to sit through adverse weather (Ronconi et al., 2015; Sarà et al., 2023; Shamoun-Baranes et al., 2017). It is conceivable that (remote) islands play an underappreciated role as stepping-stones or emergency stop-overs in many trans-oceanic migrations.

In this study, we aimed to better understand how long-distance migrant Eleonora’s falcons *Falco eleonorae* mitigate the risk of crossing the Indian Ocean on the way to and from their Malagasy non-breeding grounds. It’s been well established that falcons from throughout the species’ breeding range spend the non-breeding season in northern Madagascar (Kassara et al 2017), and they cross the ocean along distinct corridors in autumn and spring. More specifically, they cross the Indian ocean at its narrowest point near the Mozambique Channel (∼420 km) in opposing autumn winds (López-López et al., 2010). By contrast, they cover at least twice that distance (and often much more) by crossing directly from northern Madagascar to mainland East Africa in supportive spring winds (Gschweng et al., 2008; López-López et al., 2010, Mellone et al 2013, Hadjikyriakou et al 2020, Vansteelant et al 2021). We also know that individual falcons exhibit highly flexible spring routes over the Indian Ocean depending on annual variations in wind conditions (Mellone et al., 2011). However, it remains unclear to what extent the supportive winds in spring compensate for the additional distance flown over the ocean in spring compared to autumn, and how falcons mitigate the risk of ocean-crossings by adjusting their flight times and speeds (i.e. air speeds) to prevailing winds. Moreover, there are a number of small islands, archipelagos, and atolls along the way that may serve as stepping stones (Nourani et al., 2018) or emergency stop-over sites (including Comoros, Glorioso Islands and Aldabra group, Fig.1). There is anecdotal evidence thatfalcons sometimes stop-over (tracking) or perish on these islands (ring recoveries) (Gschweng et al., 2008). However, it is unknown how regularly Eleonora’s falcons use islands, and if so, under what environmental conditions, for how long, and what is the function of island visits (i.e. resting, foraging or both).

**Fig. 1.**
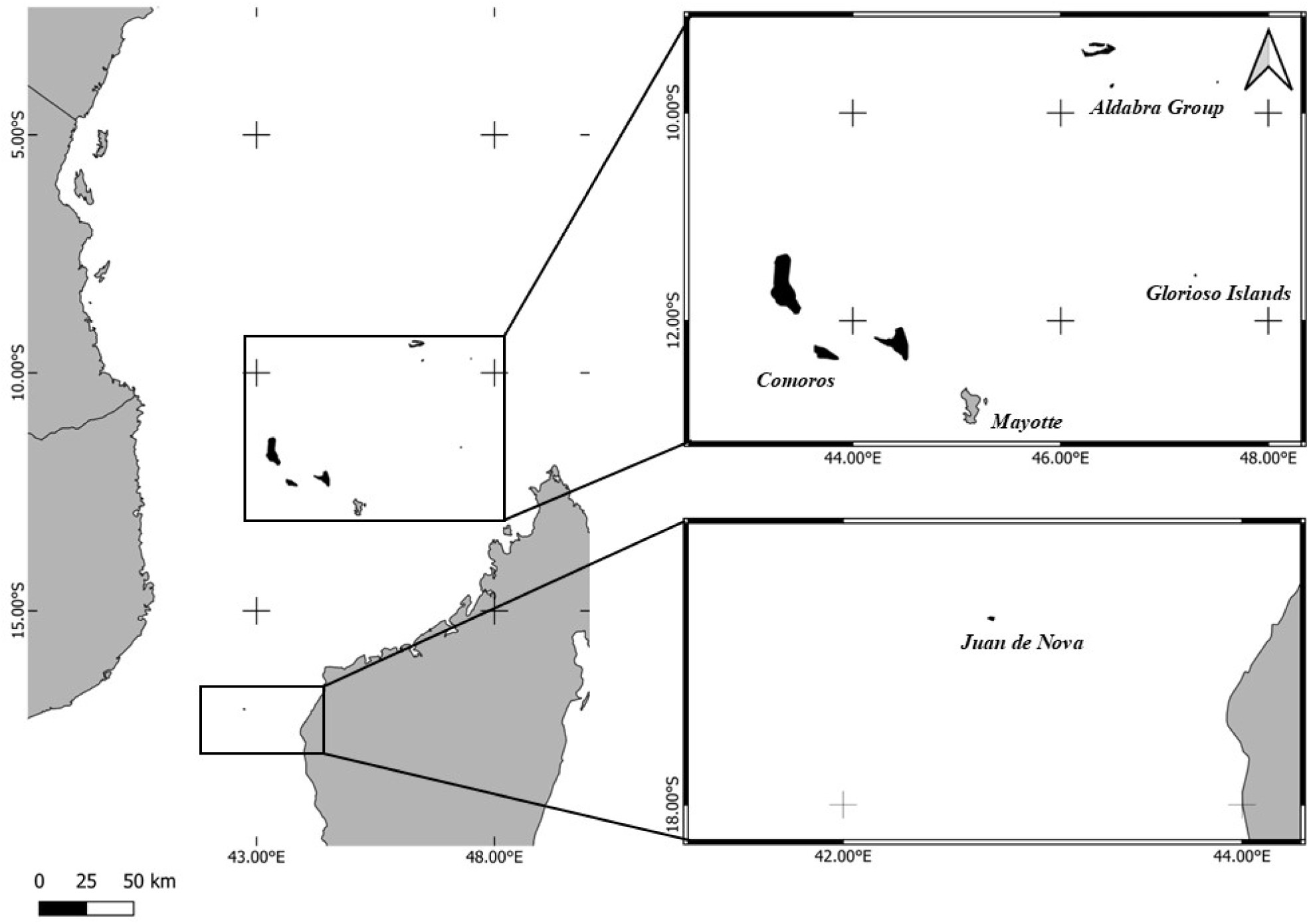
Study area in the Western Indian Ocean. Geographic extent of the study area includes the Aldabra Group, Glorioso Islands, Mayotte, the Comoros, and Juan de Nova. Insets show magnified views of the northern and southern island clusters. Shaded regions indicate surrounding landmasses, including north Madagascar and the eastern African coast.

Our study leverages high-resolution GPS-tracking data from Eleonora’s Falcons studied at the species’ westernmost population on the Canary Islands. Like conspecifics from other parts of the breeding range, falcons from our study population use distinct seasonal corridors over the Indian Ocean (Vansteelant et al 2021). We (i) quantify seasonal differences in ocean-crossing distance and duration, and the use of islands, (ii) quantify the diel timing of trans-oceanic flights (incl. timing of take-off and landings from/on maindland and islands), and (iii) assess diel activity patterns of falcons on islands. Next, we (iv) test for seasonal differences in instantaneous and cumulative wind support (average tailwind speed and overall wind displacement), and falcons’ instantaneous and cumulative flight effort over the ocean (i.e. average air speeds and air distances), and finally, (v) test whether wind conditions experienced by falcons over the ocean affect the likelihood of them using islands in either season.

We know from previous research that falcons are more likely to encounter islands along their spring ocean-crossing corridor than over the Mozambique channel in autumn (Vansteelant et al 2021, Fig.1), and we therefore expect that a greater proportion of individuals used islands during spring than autumn crossings. We also know that falcons receive greater wind support during spring ocean-crossings, and here we epect that this allows falcons to cross at lower airspeeds, i.e. to reduce instantaneous flight cost compared to autumn. At the same time, while wind support should compensate for much of the additional ocean-crossing distance in spring, we expect falcons still make a greater cumulative flight effort (i.e. air distance) over the ocean in spring, and that they use islands as emergency stop-overs when encountering relatively weak wind support.

## Methods

### Study area

This study investigates the migratory patterns of Eleonora’s falcons across the Indian Ocean and Mozambique Channel, including the potential role of islands located 100–400+ km offshore from East Africa or Madagascar. Along these routes, several island groups - including the Seychelles (particularly the Aldabra Group), the Glorioso Islands, Mayotte, the Comoros (Grande Comore, Mohéli, and Anjouan), and Juan de Nova - are positioned such that they could, in theory, function as stepping stones or emergency stop-over sites for migrating falcons in one or both seasons (Fig. 1).

### Falcon tracking

We tracked Eleonora’s falcons from the species’ westernmost breeding population on the Canary Islands. Adult falcons were caught at the nest or at a drinking pool on the islet of Alegranza, which holds about 127 breeding pairs representing just under half of the Canarian population (Gangoso et al., 2020, 2015). UvA-BiTS GPS loggers, weighing 7.5 g, were attached to falcons using a Teflon body harness (Gangoso et al., 2020; Viana et al. 2016). Data was downloaded through a locally constructed network of antennas, and we obtained migration data for 19 adult individuals that returned to Alegranza after completing their migration cycle to and from Madagascar. More details on catching and tagging procedures, return and data retrieval rates are provided by Vansteelant et al. (2021). The devices recorded positions at varying intervals depending on battery levels, and for the ocean-crossing flights the temporal resolution of the data varied from 0.1-10 min intervals for 99% of all data points.

### Identifying ocean-crossings and island visits

We identified the start and end points of crossings based on their geographic location. More specifically, we selected the last/first location on Tanzania, Kenya or Mozambique as the start of autumn and the end of spring crossings, respectively. Analogously, we selected the first/last point on Madagascar as the end of autumn and the start of spring crossings, respectively. Islands near (i.e. <50km from) the East African and Malagasy shores were included in this definition. In total, our dataset comprised 83 sea-crossing events by 19 individuals across 10 years (2012-2022), including 42 autumn and 41 spring sea-crossing events. For our analysis of the use of islands as stepping-stones or stop-overs, we focus on those islands and atolls that are situated at least 100km (Juan de Nova) and in most cases >250km from the nearest major landmass.

To ensure even the smallest islands were included, we created a custom shapefile by tracing island/atoll contours in Google Earth v7.3.6.1020 (imagery March 12, 2024) and identified every unique island visit.

### Estimating wind support

All GPS positions were annotated with wind data from the NOAA-NCEP Reanalysis II model (resolution 6h and 2.5°x 2.5°; Kalnay et al 1996, Kemp et al 2010). Based on the falcons recorded flight altitudes, we decided to use the u- and v-wind components at the 925 mb pressure level, corresponding to approximately 750 meters above sea level. Using basic trigonometry we then converted the u- and v-wind components to wind speed and direction, calculated the speed of tail-/headwind and sidewinds relative to the birds’ travel directions (Vansteelant et al., 2021), and finally the birds’ airspeed from each GPS-position to the next. In addition, to visualize seasonal wind fields over the Indian Ocean, we extracted u- and v-wind estimates for each model run during the autumn and spring ocean-crossing periods, and calculated the average wind strength and speed for every node in the model grid. The reanalysis models have a much coarser temporal resolution than the tracking data, and are known to be less accurate in parts of the global south (Kalnay et al 1996). However, we expect that the reanalysis model captures major seasonal patterns in prevailing, synoptic winds, as well as day-to-day differences in wind strength and direction over the Indian Ocean and Mozambique Channel (Vansteelant et al., 2021).

### Quantifying seasonal ocean-crossing metrics

After extracting ocean-crossings and annotating them with island and wind data, we calculated several behavioural and wind support metrics for each crossing: the total duration of each crossing (including island visits), the total amount of time spent on islands, the number of islands used, the number of nocturnal and diurnal flight hours, the average ground speed, tail-/headwind strength and airspeed during each crossing (i.e. excl. time spent on islands), the ground distance, cumulative wind-induced displacement over the ocean (with negative values indicating cases where falcons were ‘pushed back’ by headwinds), and the cumulative air distance flown over the ocean. Airspeed and air distance can, respectively, be considered as proxies for the falcons’ instantaneous and cumulative energy expenditure in active flapping flight over the ocean (Pennycuick, 2008; Liechti, 2006).

### Diel timing of ocean-crossings and activity patterns on islands

We assessed diel timing of ocean-crossings by calculating at what ‘solar time’ falcons departed or arrived at either end of the ocean-crossing, and during island visits. We quantified peaks in departure and arrival frequencies relative to (civil) sunrise and sunset, and assessed diel activity during island visits using ground speeds to explore whether falcons were using islands to rest or forage during night and day.

### Statistical approach

To evaluate the significance of seasonal differences in aforementioned flight, island use and wind support metrics we utilized a Generalized Linear Mixed Modeling approach (GLMM). We included a fixed effect for season, to estimate the average difference between autumn and spring estimates, and random intercepts for individual birds to account for repeated measurements of the same individuals. Most of the variables satisfied the assumption of normally distributed residuals.

For the variable “Island Count”, an integer variable, residuals best fitted a Poisson distribution. In cases where the estimation of random individual ilntercepts in mixed models resulted in singular fits we instead adopted generalized linear models (GLM).

Next, we aimed to analyze how departure location (specifically latitude) and wind conditions experienced over the ocean affected the probability of a falcon using an island. We quantified wind support during the “first half” of the ocean-crossing because falcons typically encounter islands around the mid-way point of seasonal ocean-crossings. We defined the ‘first half’ as the moment of initial take-off until (i) the first fix recorded over an island or, (ii) in cases where falcons did not pass over an island, a hand-drawn boundary representing the approximate half-way point along the autumn and spring ocean-crossing corridors, respectively. Following this segmentation we could calculate the mean head-/tailwind that falcons experienced during the first half of each sea-crossing. We again used GLLMs to implement a logistic regression for each season separately, with the probability of using an island (0,1) as response variable, the average wind support during the first half of the crossing and departure latitude as fixed effects, and allowing for randomly varying intercepts between individuals to account for non-independence among repeated measurements of the same individuals. Finally, to help assess the way in which falcons use islands, we used GLMMs with a beta residual distribution to test whether the proportion of time spent resting was correlated with the time spent on the island at night or by day and the total duration of island visits. In all cases where the inclusion of random individual intercepts in GLMMS resulted in singular fits, we corroborated effects using GLMs instead.

### Software and Tools

All statistical analyses and data visualization were conducted in R version 4.3.1 (R Core Team, 2023), except for Fig.1 which was made in QGIS (v.3.30.2).

## Results

### General description of seasonal ocean-crossings

Falcons started their autumn ocean-crossings on 14^th^ November ± 5.3 days, starting from the coast of Mozambique or Tanzania to cross the Mozambique Channel near its narrowest point (Fig.2A). The ocean-crossing corridor was bordered by the islands of Mayotte and the Comoros to the north. Falcons typically experienced headwinds during autumn crossings, though some birds that crossed south of the narrowest point of the Channel experienced weak-moderate tailwinds (Fig.2A, note north-south gradient in track colours). By crossing south of Mayotte and the Comoros, the falcons effectively avoided an area with much stronger potential headwinds north of the islands (Fig. 2A: note longer wind arrows and darker background shading).

**Fig. 2.**
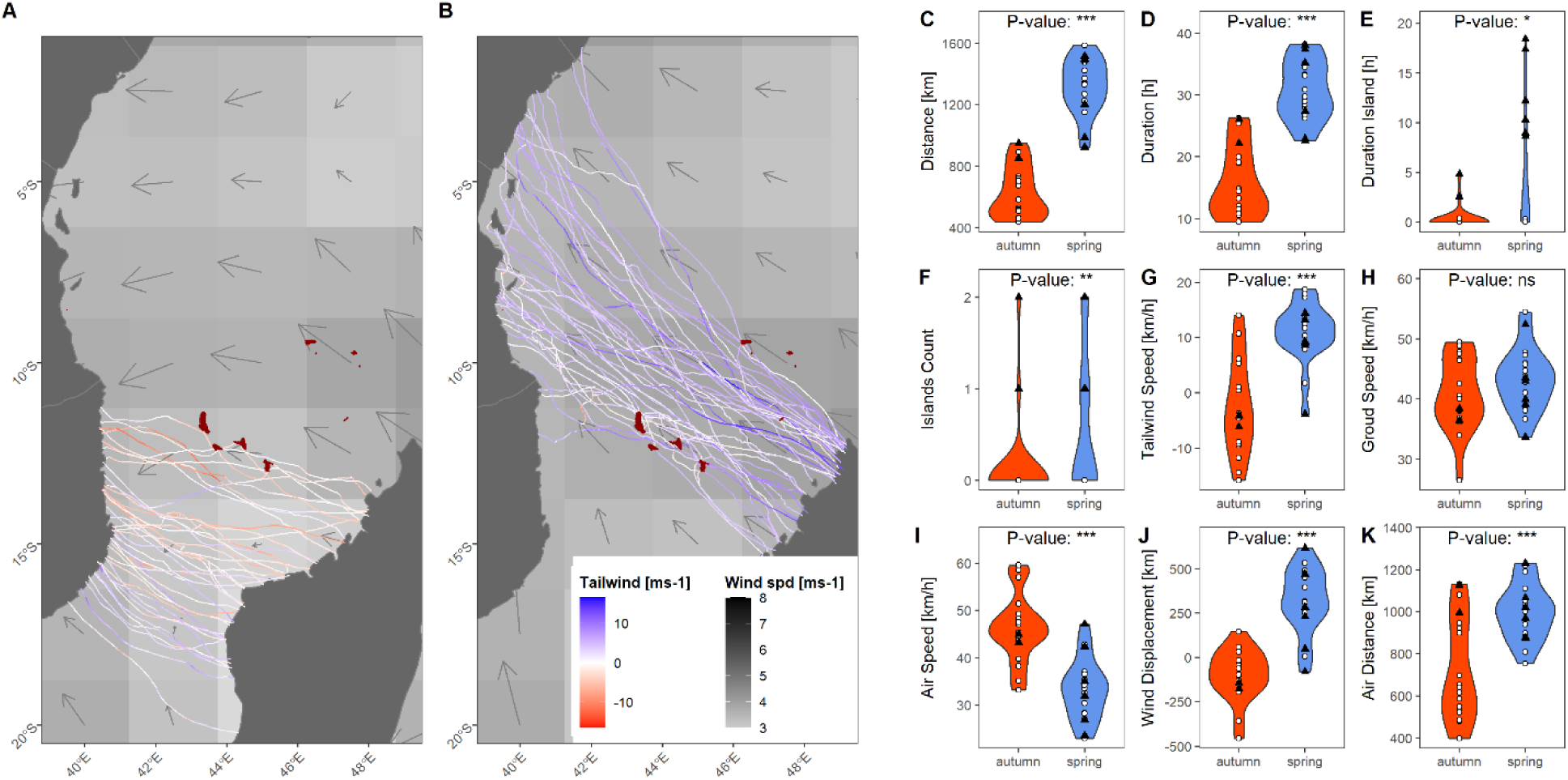
Seasonal comparison of Eleonora’s falcons’ ocean-crossings between East Africa and Madagascar. (A,B) Ocean-crossing trajectories coloured according to tailwind support along the falcons’ realized travel direction (reds = headwinds, blues = tailwinds). Background shading and arrows indicate average wind conditions over the ocean in 2.5° x 2.5° cells, with darker shading and longer arrows indicating stronger winds. Remote islands situated within the falcons’ seasonal crossing corridors are shown in dark red. (C-K) Violin plots showing the seasonal distribution of ocean-crossing parameters, based on the first recorded crossing of each individual: (C) total distance flown over the ocean, (D) total crossing duration, (E) time spent on islands, number of islands used, (G) average tail-/headwind strength along the falcons’ realized tracks, (H) average ground speeds, (I) average air speeds (J) cumulative wind displacement, and (K) total air distance flown over the ocean. Points indicate values for individual crossings, with white dots indicating crossings on which falcons did not use any island, and black triangles indicating crossings on which falcons used at least one island. Labels indicate significance of seasonal differences according to pairwise t-tests (ns: not significant, *: 0.05 < P < 0.01; **: P < 0.01 ; ***: P < 0.001).

Falcons started their spring ocean-crossings on 10^th^ April ± 7.5 days, departing from northern Madagascar in a northwestern direction to mainland East Africa (Fig.2B). The transoceanic spring corridor was bordered by the islands of Mayotte and the Comoros to the south, and the Seychelles’ Aldabra group to the north (Fig. 2B). In general, falcons seemed to enjoy moderate-strong tailwinds (Fig.2B, note strong dominance of blue coloured tracks). The spring ocean-crossing corridor seemed to overlap with the area where prevailing (south-) easterlies were strongest (Fig.2B, note background shading is darkest where falcons cross in spring).

### Seasonal ocean-crossing distance, duration and island use

Autumn ocean-crossings spanned a geographical distance of 400-1000 km (Fig.2C, Table S1), which falcons typically completed in less than a day (15.3 ± 5.4 h, Fig.2D, Table S1). Spring crossings were significantly longer at 800-1600 km (Fig.2C, Table S1), and typically took twice as long - usually more than a day (31.0 ± 5.2 h, Fig.2D, Table S1) to complete. The difference in crossing duration was associated with a greater amount of time spent on greater number of islands in spring (Fig. 2E-F, Table S1). This was due to a greater proportion of individuals using islands in spring (Table 1) as well as island visits taking longer in spring (Tables 1-2). Island use was not the norm in either season, as >60% of the ocean-crossings were completed in non-stop flight.

**Table 1.**
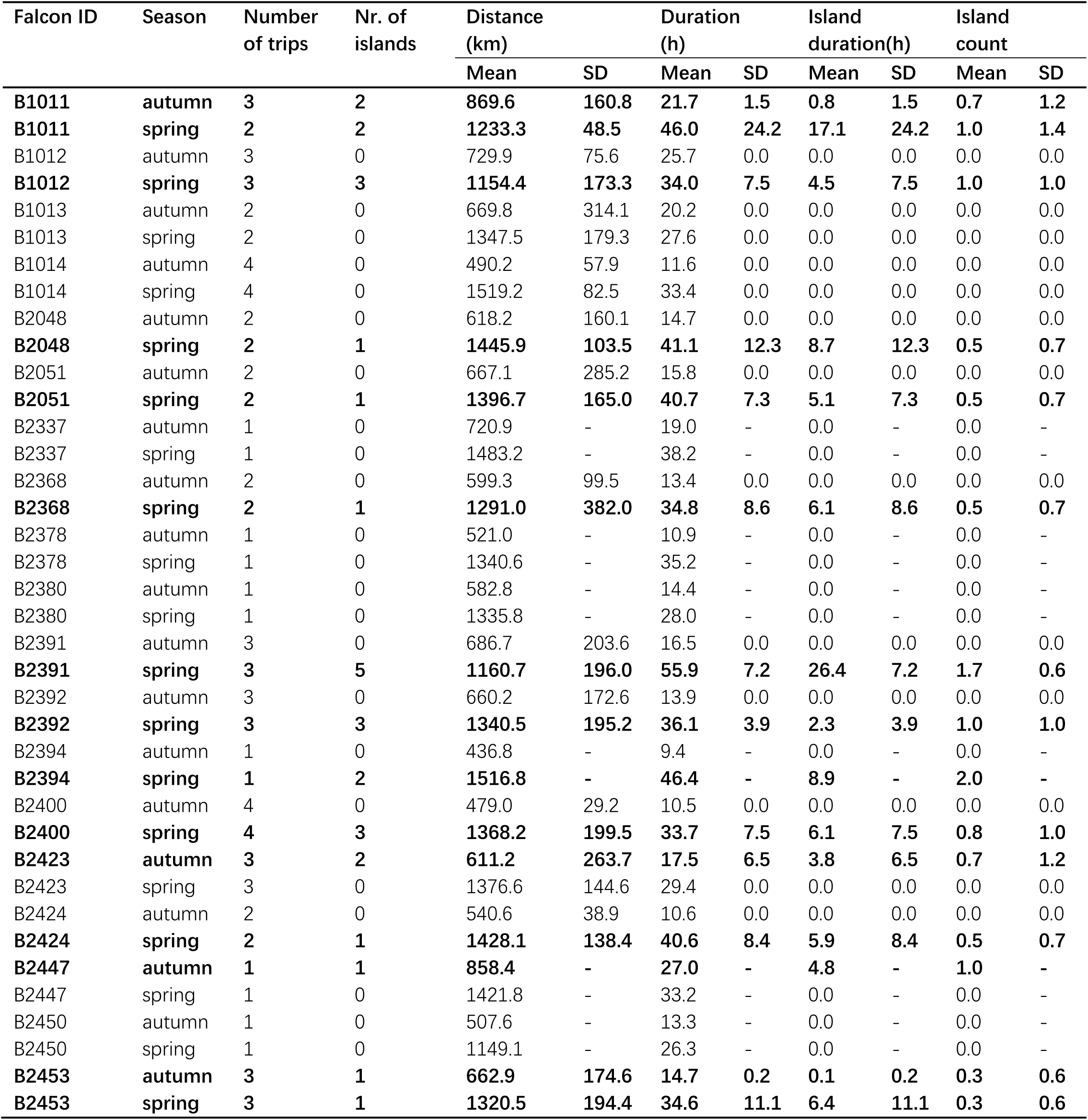
Summary statistics including seasonal sample sizes, ocean-crossing distance and duration, number of islands used, and time spent on islands, corresponding to Fig.2C-F. Bold rows indicate individuals that used at least one island in one of their recorded journeys.

| Falcon ID | Season | Number of trips | Nr. of islands | Distance (km) |  | Duration (h) |  | Island duration(h) |  | Island count |  |
| --- | --- | --- | --- | --- | --- | --- | --- | --- | --- | --- | --- |
|  |  |  |  | Mean | SD | Mean | SD | Mean | SD | Mean | SD |
| **B1011** | **autumn** | **3** | **2** | **869.6** | **160.8** | **21.7** | **1.5** | **0.8** | **1.5** | **0.7** | **1.2** |
| **B1011** | **spring** | **2** | **2** | **1233.3** | **48.5** | **46.0** | **24.2** | **17.1** | **24.2** | **1.0** | **1.4** |
| B1012 | autumn | 3 | 0 | 729.9 | 75.6 | 25.7 | 0.0 | 0.0 | 0.0 | 0.0 | 0.0 |
| **B1012** | **spring** | **3** | **3** | **1154.4** | **173.3** | **34.0** | **7.5** | **4.5** | **7.5** | **1.0** | **1.0** |
| B1013 | autumn | 2 | 0 | 669.8 | 314.1 | 20.2 | 0.0 | 0.0 | 0.0 | 0.0 | 0.0 |
| B1013 | spring | 2 | 0 | 1347.5 | 179.3 | 27.6 | 0.0 | 0.0 | 0.0 | 0.0 | 0.0 |
| B1014 | autumn | 4 | 0 | 490.2 | 57.9 | 11.6 | 0.0 | 0.0 | 0.0 | 0.0 | 0.0 |
| B1014 | spring | 4 | 0 | 1519.2 | 82.5 | 33.4 | 0.0 | 0.0 | 0.0 | 0.0 | 0.0 |
| B2048 | autumn | 2 | 0 | 618.2 | 160.1 | 14.7 | 0.0 | 0.0 | 0.0 | 0.0 | 0.0 |
| **B2048** | **spring** | **2** | **1** | **1445.9** | **103.5** | **41.1** | **12.3** | **8.7** | **12.3** | **0.5** | **0.7** |
| B2051 | autumn | 2 | 0 | 667.1 | 285.2 | 15.8 | 0.0 | 0.0 | 0.0 | 0.0 | 0.0 |
| **B2051** | **spring** | **2** | **1** | **1396.7** | **165.0** | **40.7** | **7.3** | **5.1** | **7.3** | **0.5** | **0.7** |
| B2337 | autumn | 1 | 0 | 720.9 | - | 19.0 | - | 0.0 | - | 0.0 | - |
| B2337 | spring | 1 | 0 | 1483.2 | - | 38.2 | - | 0.0 | - | 0.0 | - |
| B2368 | autumn | 2 | 0 | 599.3 | 99.5 | 13.4 | 0.0 | 0.0 | 0.0 | 0.0 | 0.0 |
| **B2368** | **spring** | **2** | **1** | **1291.0** | **382.0** | **34.8** | **8.6** | **6.1** | **8.6** | **0.5** | **0.7** |
| B2378 | autumn | 1 | 0 | 521.0 | - | 10.9 | - | 0.0 | - | 0.0 | - |
| B2378 | spring | 1 | 0 | 1340.6 | - | 35.2 | - | 0.0 | - | 0.0 | - |
| B2380 | autumn | 1 | 0 | 582.8 | - | 14.4 | - | 0.0 | - | 0.0 | - |
| B2380 | spring | 1 | 0 | 1335.8 | - | 28.0 | - | 0.0 | - | 0.0 | - |
| B2391 | autumn | 3 | 0 | 686.7 | 203.6 | 16.5 | 0.0 | 0.0 | 0.0 | 0.0 | 0.0 |
| **B2391** | **spring** | **3** | **5** | **1160.7** | **196.0** | **55.9** | **7.2** | **26.4** | **7.2** | **1.7** | **0.6** |
| B2392 | autumn | 3 | 0 | 660.2 | 172.6 | 13.9 | 0.0 | 0.0 | 0.0 | 0.0 | 0.0 |
| **B2392** | **spring** | **3** | **3** | **1340.5** | **195.2** | **36.1** | **3.9** | **2.3** | **3.9** | **1.0** | **1.0** |
| B2394 | autumn | 1 | 0 | 436.8 | - | 9.4 | - | 0.0 | - | 0.0 | - |
| **B2394** | **spring** | **1** | **2** | **1516.8** | **-** | **46.4** | **-** | **8.9** | **-** | **2.0** | **-** |
| B2400 | autumn | 4 | 0 | 479.0 | 29.2 | 10.5 | 0.0 | 0.0 | 0.0 | 0.0 | 0.0 |
| **B2400** | **spring** | **4** | **3** | **1368.2** | **199.5** | **33.7** | **7.5** | **6.1** | **7.5** | **0.8** | **1.0** |
| **B2423** | **autumn** | **3** | **2** | **611.2** | **263.7** | **17.5** | **6.5** | **3.8** | **6.5** | **0.7** | **1.2** |
| B2423 | spring | 3 | 0 | 1376.6 | 144.6 | 29.4 | 0.0 | 0.0 | 0.0 | 0.0 | 0.0 |
| B2424 | autumn | 2 | 0 | 540.6 | 38.9 | 10.6 | 0.0 | 0.0 | 0.0 | 0.0 | 0.0 |
| **B2424** | **spring** | **2** | **1** | **1428.1** | **138.4** | **40.6** | **8.4** | **5.9** | **8.4** | **0.5** | **0.7** |
| **B2447** | **autumn** | **1** | **1** | **858.4** | **-** | **27.0** | **-** | **4.8** | **-** | **1.0** | **-** |
| B2447 | spring | 1 | 0 | 1421.8 | - | 33.2 | - | 0.0 | - | 0.0 | - |
| B2450 | autumn | 1 | 0 | 507.6 | - | 13.3 | - | 0.0 | - | 0.0 | - |
| B2450 | spring | 1 | 0 | 1149.1 | - | 26.3 | - | 0.0 | - | 0.0 | - |
| **B2453** | **autumn** | **3** | **1** | **662.9** | **174.6** | **14.7** | **0.2** | **0.1** | **0.2** | **0.3** | **0.6** |
| **B2453** | **spring** | **3** | **1** | **1320.5** | **194.4** | **34.6** | **11.1** | **6.4** | **11.1** | **0.3** | **0.6** |

**Table 2.** Island use by Eleonora’s Falcons during autumn and spring ocean-crossing.

| Season | Island | Mean island stopover duration (h) | Nr. of individuals | Nr. of trips |
| --- | --- | --- | --- | --- |
| autumn | Anjouan | 0.11 | 3 | 3 |
| autumn | Mayotte | 3.46 | 2 | 2 |
| autumn | Mohéli | 10.38 | 1 | 1 |
| spring | Aldabra | 10.27 | 3 | 3 |
| spring | Anjouan | 18.07 | 1 | 1 |
| spring | Assomption Grande | 0.03 | 1 | 1 |
| spring | Comore Grande | 11.23 | 5 | 6 |
| spring | Glorieuse | 8.54 | 2 | 2 |
| spring | Mayotte | 11.37 | 4 | 6 |
| spring | Mohéli | 8.62 | 4 | 4 |

However, 68% of the falcons did use an island in at least one of the years they were tracked, and the frequency of island use was much higher in spring (57% of falcons) compared to autumn (15% of falcons, Table 1). The islands used by most individuals were Comore Grande (26% of falcons) and Moheli (21% of falcons) in the Comoros, and the island of Mayotte (21% of falcons) (Table 2).

### Diel timing of ocean-crossings and activity on islands

Ocean-crossings involved substantial sections of nocturnal flight in both autumn (Fig.3A) and spring (Fig.3B). On average, falcons flew significantly more hours at night during the much longer spring crossings (Fig.3C, Table S2). However, there was no significant difference in the average proportion of nocturnal flight time, and on average <50% of the total crossing was conducted at night in both seasons (Fig.3D, Table S2). There was, however, much greater variation in the amount and proportion of nocturnal flight hours during autumn than in spring, with the vast majority of spring crossings involving 12-15h of nocturnal flight amounting to 30-50% of total crossing time (Fig.3C-D). The notable exceptions were two cases where falcons reduced nocturnal travel time to ca. 3-4h, i.e. <20% of total crossing time, by overnighting on islands (Fig. 3C-D, note both “outlier” data points are black triangles).

**Fig. 3.**
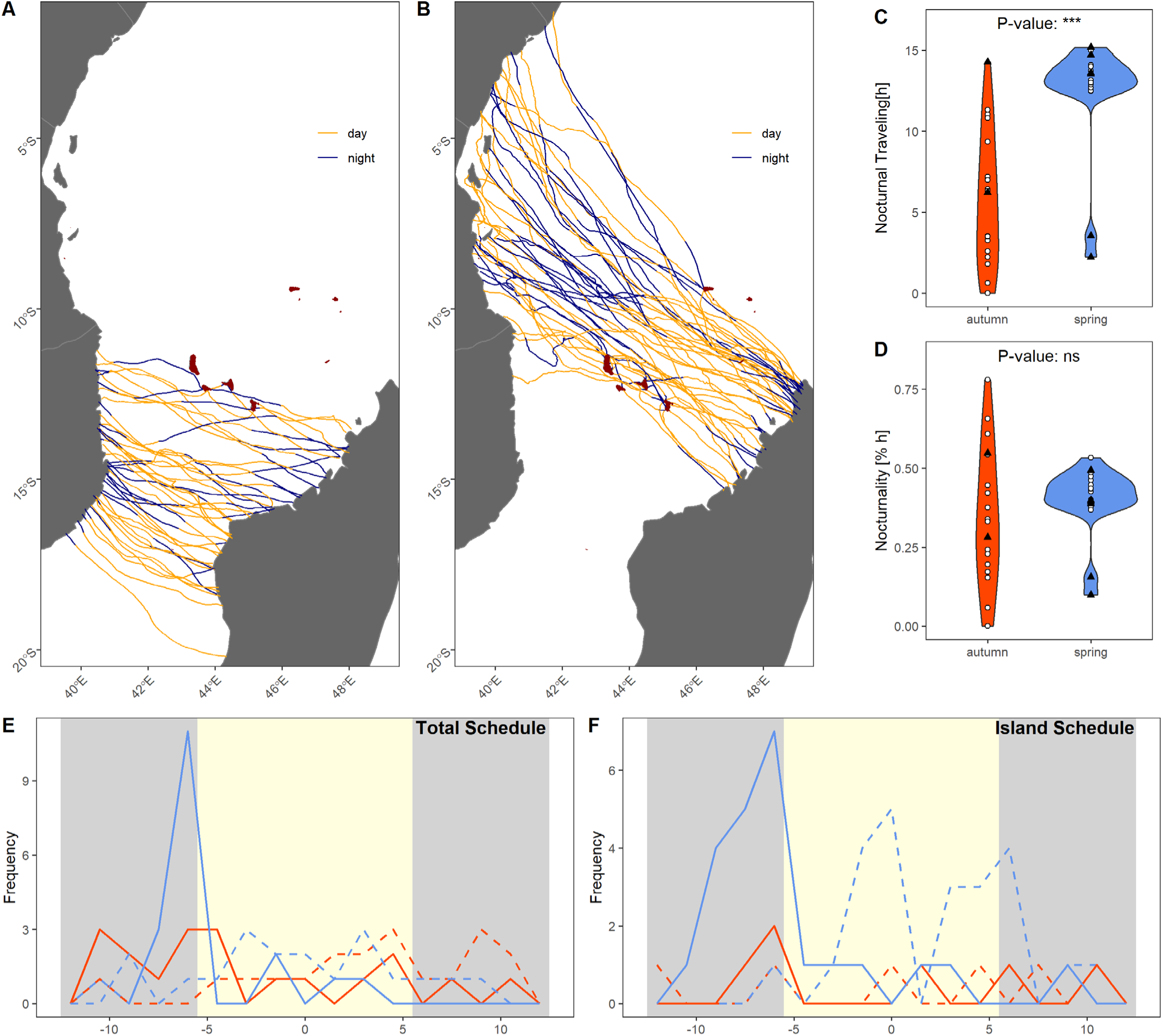
Nocturnality during ocean-crossings along with diel timing of take-offs and arrivals. (A) Autumn and (B) spring ocean-crossing routes indicating nocturnal (blue, including twilight) and diurnal (orange) flight. (C) The number of nocturnal flight hours and (D) proportion of nocturnal flight hours, with lay-out analogous to panels Fig.2C-I. (E-F) Frequency distributions of take-off times (solid lines) and arrival times (dashed lines) times (E) at either end of the ocean-crossing and (F) on islands. X-axis shows solar time (i.e. hours vs solar noon). The shading in the background indicates the average daytime period (light yellow) and night-time period (light grey) on the falcon’s ocean-crossing dates.

Initiation of ocean-crossings was highly concentrated in early dawn hours in spring, both from Madagascar (Fig.3E, solid blue line) and when departing from islands (Fig.3F). In autumn, crossings were initiated more spread out throughout the day. In both seasons, falcons completed sea-crossings (Fig.3E, dashed lines) or reached islands (Fig.3F, dashed lines) at spread out times throughout the day, and to a lesser extent at night.

Trajectory speeds recorded on islands indicate that falcons were active during the day and resting at night (Fig.S1A-D). When using an island in spring (typically in the 2^nd^ half of the day), falcons seemed to almost always stay on the island until the next morning, so that most of their time on islands was used for resting at night, (Fig.S1E-F). Earlier arrival times did not result in longer rest periods; indicating that falcons that reached islands during the day spent the rest of the daytime period patrolling or foraging rather than resting (Table S4).

### Seasonal differences in wind support and flight effort

The NOAA-NCEP Reanalysis II model indicates that, on average, falcons experienced headwinds of −1.7 ± 9.3 km/h in autumn and moderate tailwinds of 13.8 ± 9.0 km/h in spring. While we did not formally test whether falcons selected days with relatively favorable winds to initiate seasonal crossings, winds generally opposed the falcons travel direction on the great majority of days in autumn, and ocean-crossing dates were not clearly linked to rare windows of supportive winds (Fig.S2A). By contrast, prevailing winds provide moderate-strong tailwind assistance along the falcons’ mean flight direction on almost every day of the spring migration period (Fig.S2B).

Despite the strong seasonal contrast in tail-/headwinds experienced over the ocean, falcons crossed the ocean at comparable ground speeds of 42.0 ± 7.2 km/h in autumn and 43.5 ± 5.4 km/h in spring. As such, they flew at significantly higher airspeeds in autumn (47.4 ± 9.3 km/h) than in spring (31.5 ± 6.0 km/h). Integrated over the entire crossing, falcons were ‘pushed backwards’ over 93.0 ± 152.5 km by adverse autumn winds, while in spring winds falcons were ‘pushed forward’ 367.1 ± 193.4 km (Fig. 2C-E, Table S1). Consequently, the seasonal difference in ‘air distances’ was much smaller than the seasonal difference in ‘ground distances’ flown over the ocean. Nevertheless, falcons still made a significantly greater flight effort over the ocean in spring (965.2 ± 192.8 km) than in autumn (717.4 ± 263.3 km).

### Wind effects on island use

We found evidence that island use was correlated with wind conditions experienced during the first half of the ocean-crossing in spring, but not in autumn (Fig.4, Table S3). A single-effect logistic regression model did not reveal a significant effect of wind conditions on the probability of island use in either season (Fig. 4, Table S3). However, models that account for departure latitude show that the probability of island use and time spent on islands significantly increased with weaker wind support in spring (Table S3). The effect of departure latitude is due to the fact that in autumn falcons did not use the San Juan island in the Mozambique channel, and only crossings that were initiated relatively far north on the East African coast were likely to include the Mayotte and Comoros Islands. In spring, falcons departing from the southern end of the crossing corridor were much more likely to use the Mayotte and Comoros islands than those departing from the northern end, which mostly used the Aldabra group of the Seychelles (Table 2).

**Fig. 4.**
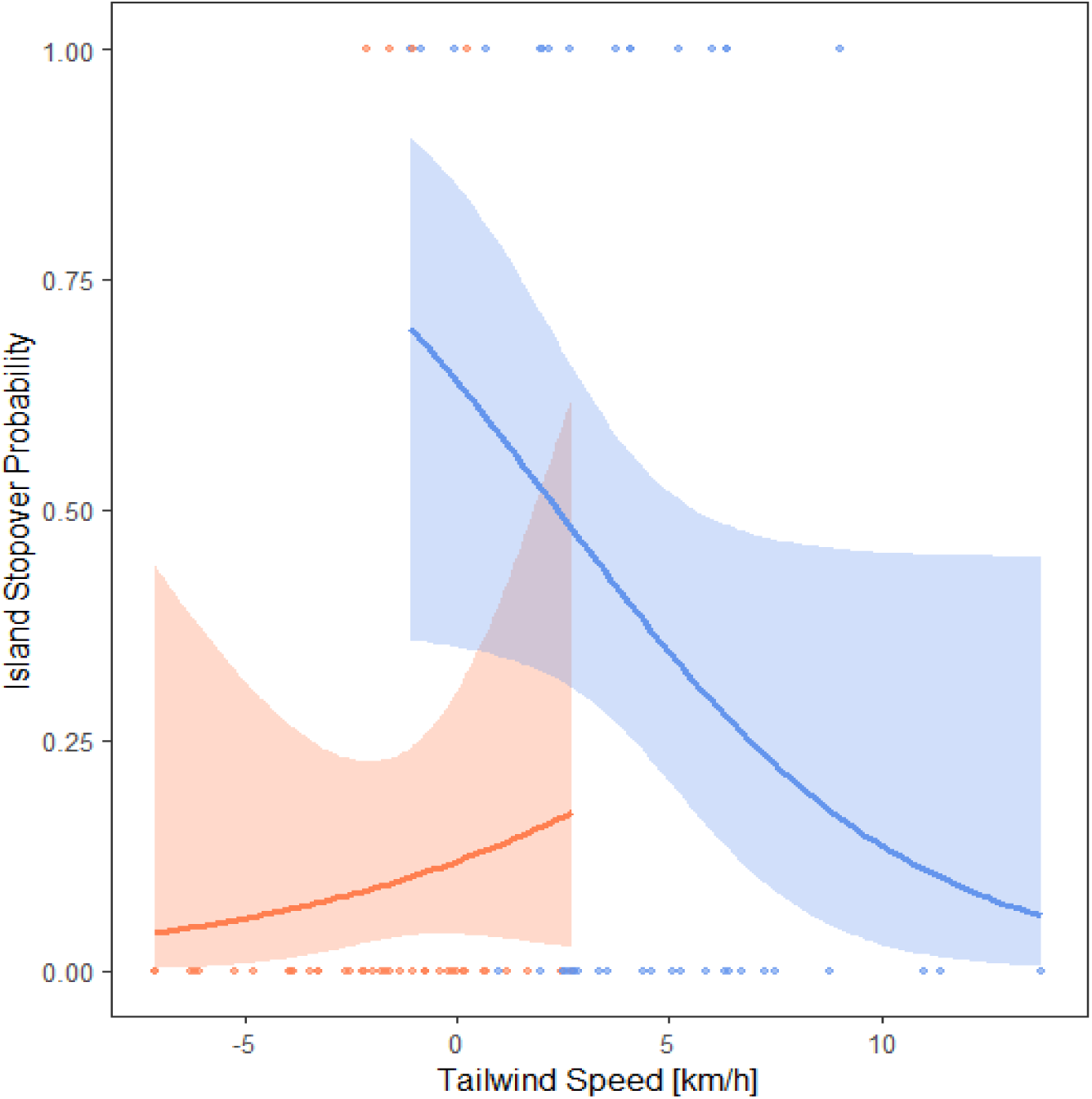
Logistic regressions for the effect of average tailwind strength during the first half of ocean-crossings on island stopover probability.

## Discussion

Our detailed analyses of high-resolution movement data of Eleonora’s falcons provides further support for the importance of prevailing winds in shaping these landbirds’ seasonal ocean-crossing corridors (Mellone et al., 2011; Nourani et al., 2021; Vansteelant et al., 2021). Moreover, our work reveals additional risk-mitigating behaviours, including wind-dependent airspeed adjustments and use of remote islands. Falcons use islands especially when they experience relatively weak tailwind support during their ca. 1000km spring ocean-crossings, so that we interpret this as an example of emergency stop-overs (cf. Shamoun-Baranes et al. 2010, 2017). Falcons appear to mitigate risk by flying at much lower air speed in supportive spring winds than in adverse autumn winds. Such airspeed adjustments allow falcons to reduce instantaneous energy expenditure, and to increase their potential flight time and range (Vansteelant, Shamoun-Baranes, McLaren, et al., 2017).

This is a sensible strategy considering that the seasonal contrast in wind support does not fully offset the seasonal difference in ocean-crossing distance and duration. Indeed, even while flying through the area with most supportive prevailing winds over the Indian Ocean in spring, and through the area with weakest headwinds in autumn, falcons still cover a 40% greater air distance over the ocean in spring than in the majority of autumn crossings. Best make such flights at a moderate marathon pace.

In general, seasonal and regional differences in the ground speeds of migratory birds are largely explained by seasonal and regional variations in wind support (Liechti, 2006; Shamoun-Baranes et al., 2017; Vansteelant et al., 2015) . When considering the entire trans-African migration, this is also true for the Eleonora’s falcons we studied here, which were previously shown to complete their ca. 20% longer spring migrations in the same amount of flight hours as in autumn, thanks largely to falcons attaining higher ground speeds in supportive winds in spring (Vansteelant et al., 2021). In this sense, it seems surprising that the large seasonal difference in wind support over the ocean does not result in seasonal differences in ocean-crossing ground speeds. However, it has also been established that the relationship between tailwind support and ground speeds varies across different regions of the falcons’ trans-African flyways (López-López et al., 2014; Vansteelant et al., 2021). The ground speeds they achieve over the ocean are the highest they achieve anywhere in the flyway, including the autumn desert crossing where falcons enjoy even greater tailwind support than over the Indian Ocean in spring (Vansteelant et al., 2021). This can probably be explained by the consistent use of flapping flight over the ocean (Nourani et al. 2021), as opposed to frequent soaring-gliding flight over the desert. Moreover, our results enforce the notion that regional and seasonal variation in falcons’ ground speeds are at least partly due to falcons adjusting airspeeds between energy- and time-minimizing optima depending on landscape and atmospheric contexts (Harel et al., 2016; Hedenstrom & Alerstam, 1995; Vansteelant, Shamoun-Baranes, McLaren, et al., 2017).

Migratory birds can make adaptive use of wind conditions in other ways we have not tested here, for example by tuning the timing and altitude of flight to temporal and altitudinal variation in wind support (Shamoun-Baranes et al., 2017; Manola et al., 2020, Norevik et al 2023). We know from previous work that the falcons we studied here occasionally stop-over for 1-5 days on the East

African coast before initiating the ocean-crossing in autumn (Vansteelant et al., 2021). Furthermore, there are three cases of aborted ocean-crossings in our data (pers. obs.) and one in the study of Gschweng et al. (2008). This indicates that falcons sometimes do postpone or abort crossings in particularly adverse winds (Meyer et al., 2000; Santos et al., 2020). Nevertheless, our exploration of daily wind conditions in the falcons’ seasonal departure areas suggests limited opportunities for falcons to select windows of supportive wind to cross the Mozambique channel in autumn, and little need for falcons to avoid windows of adverse winds in spring.

While previous tracking studies yielded anecdotal evidence of island use by ocean-crossing falcons (Gschweng et al., 2008; Hadjikyriakou et al., 2020), our analyses reveal that the use of remote islands is a common, although irregular and wind-dependent behaviour among experienced Eleonora’s falcons. The most used islands were Comore Grande (26% of falcons) and Moheli (21% of falcons) in the Comoros and the island of Mayotte (21% of falcons). These islands are situated in an area with relatively strong adverse winds for falcons in autumn, and with relatively weak wind support in spring. However, not all falcons that past close to these islands effectively used them, and we only found a negative effect of wind support on island use in spring. Falcons also tended to stay on these islands longer in spring, typically extending their stay through the night to depart the next morning. This is consistent with the strong tendency of Eleonora’s falcons to initiate sea-crossings in the early hours of dawn from Madagascar in spring. Falcons generally show a strong preference for diurnal migration, extending flights through the night mostly over inhospitable barriers (López-López et al., 2010; Lopez-Ricaurte et al., 2021; Strandberg et al., 2009; Vansteelant et al., 2021). The apparent tendency of Eleonora’s falcons to complete as much of the Indian Ocean crossing as possible by day in spring could be due to the fact that winds tend to be weaker, while turbulence tends to be stronger over the ocean during the night than during the day (Nourani et al., 2021). Therefore, Eleonora’s falcons might enjoy slightly greater wind support and less turbulent conditions by maximising diurnal flight time in spring. Conversely, it is striking how often falcons complete the bulk of the autumn-crossing at night, considering they could complete most of their ca. 12h autumn crossing by day if they were to initiate crossings more consistently at dawn. Weaker headwinds and greater atmospheric lift under nighttime conditions could be a reason why some falcons conduct more than half of the Mozambique channel crossing at night in adverse autumn wind fields.

Overall, our results suggest that remote islands provide emergency stop-over opportunities to falcons crossing the ocean in spring, with a little more than half of birds using an island at least once during one-four years of consecutive tracking. It is important to consider we only obtained ocean-crossing data for successful adult migrants, and it is still unclear whether mortality might be associated with inclement weather events over the Indian Ocean. Therefore, it also remains unclear to what extent these stop-over opportunities improve the survival and lifetime fitness of our study species. However, we are struck by how clearly the southern and northern limit of the falcons’ spring corridor, and the northern limit of their autumn corridor, were aligned with archipelagos in the Indian Ocean. Could it be that the falcons’ seasonal ocean-crossing corridors are shaped in part by the availability of remote islands, in addition to the seasonally prevailing winds? One way to clarify this would be to simulate viable and energy- or time-optimal routes in terms of wind support. If islands do not influence the seasonal configuration of ocean-crossing corridors, then such simulations would produce much broader viable and optimal migration corridors than we observed here.

While it was previously established that our study species displays high flexibility (and low individual repeatability) in migratory route choice and timing over the Indian Ocean (Mellone et al., 2011; Vansteelant et al., 2023), we now revealed island use and flight speed adjustments as two additional highly flexible, risk-mitigating aspects of ocean-crossing behaviour. Although some research suggests selective dissapearance of individuals with less efficient navigational behaviours (Wynn et al., 2025),there is a growing body of evidence suggesting that such behaviours can be acquired through experiential and social learning in long-lived birds (Aikens et al., 2024; Brønnvik et al., 2024; Campioni et al., 2020; Sergio et al., 2022). Tracking of juvenile Eleonora’s falcons has so far revealed that they respond differently to wind conditions than adults (López-López et al., 2014), and that they often cross the Mozambique channel later and less efficiently than adults in autumn (Gschweng et al., 2008). In fact, some juveniles do not reach the Malagasy non-breeding grounds at all on their first migration, spending their first non-breeding season on mainland Africa instead (Gschweng et al., 2008; Mellone et al., 2013). It remains to be seen whether such individuals can learn their way to Madagascar, and the ocean-crossing skills this requires, later in life.

## Acknowledgements

We are grateful to J. J. Moreno, M. de la Riva, M. Majem, and P. Gustems for their invaluable assistance with fieldwork. We extend our appreciation to the Jordán and López Arias families, owners of Alegranza and Montaña Clara islets, for providing work permits and logistical support. We thank the UvA-BiTS team for the development and maintenance of the e-Ecology infrastructure, supported by LifeWatch, and for providing access to the Dutch national e-infrastructure through SURF Cooperative. Our tracking studies were facilitated by this advanced technology, and we are particularly grateful to Willem Bouten for his excellent advice on GPS-logger programming and data collection.

## Data availability statement

Raw tracking data are stored in the UvA-BiTS database (www.uva-bits.nl). Pre-processed data used in this study are available from the Dryad Digital Repository (XXX [data will be published upon acceptance]). The code for reproducing our analyses is available in the Github repository of XX (<u>XXX [name and link will be added upon acceptance]</u>). This repository automatically installs all required packages and includes detailed instructions for reproducing the analyses and obtaining open access third-party data.

## Ethical statement

The capture and tagging of falcons were conducted under permits issued by the Dirección General de Protección de la Naturaleza (Viceconsejería de Medio Ambiente) of the Government of the Canary Islands (permit numbers: 2014/2224, 2015/3835, 2017/6829, 2020/10521, and 35/2023-00526075318). The authors declare that there are no conflicts of interest related to this study.

## Author contributions

**Meixu Chen (Corresponding author):** Conceptualization (equal); Formal analysis (lead); Methodology (equal); Visualization (equal); Writing – original draft (lead); Writing – review and editing (equal).

**Duarte S. Viana**: Data acquisition and curation (supporting); Writing – review and editing (equal).

**Jordi Figuerola:** Funding acquisition (equal); Project administration (lead); Writing – original draft (supporting); Writing – review and editing (equal).

**Laura Gangoso:** Conceptualization (equal); Investigation (lead); Data curation (lead); Formal analysis (supporting); Methodology (equal); Supervision (equal); Funding acquisition (lead); Writing – original draft (supporting); Writing – review and editing (equal).

**Wouter M. G. Vansteelant:** Conceptualization (equal); Formal analysis (supporting); Supervision (lead); Methodology (equal); Visualization (equal); Writing – original draft (supporting); Writing – review and editing (equal).

