## supplementaryfigures for "Falcons use wind assistance and remote islands to mitigate risk during ocean-crossings"

### SUPPLEMENTARY FIGURES

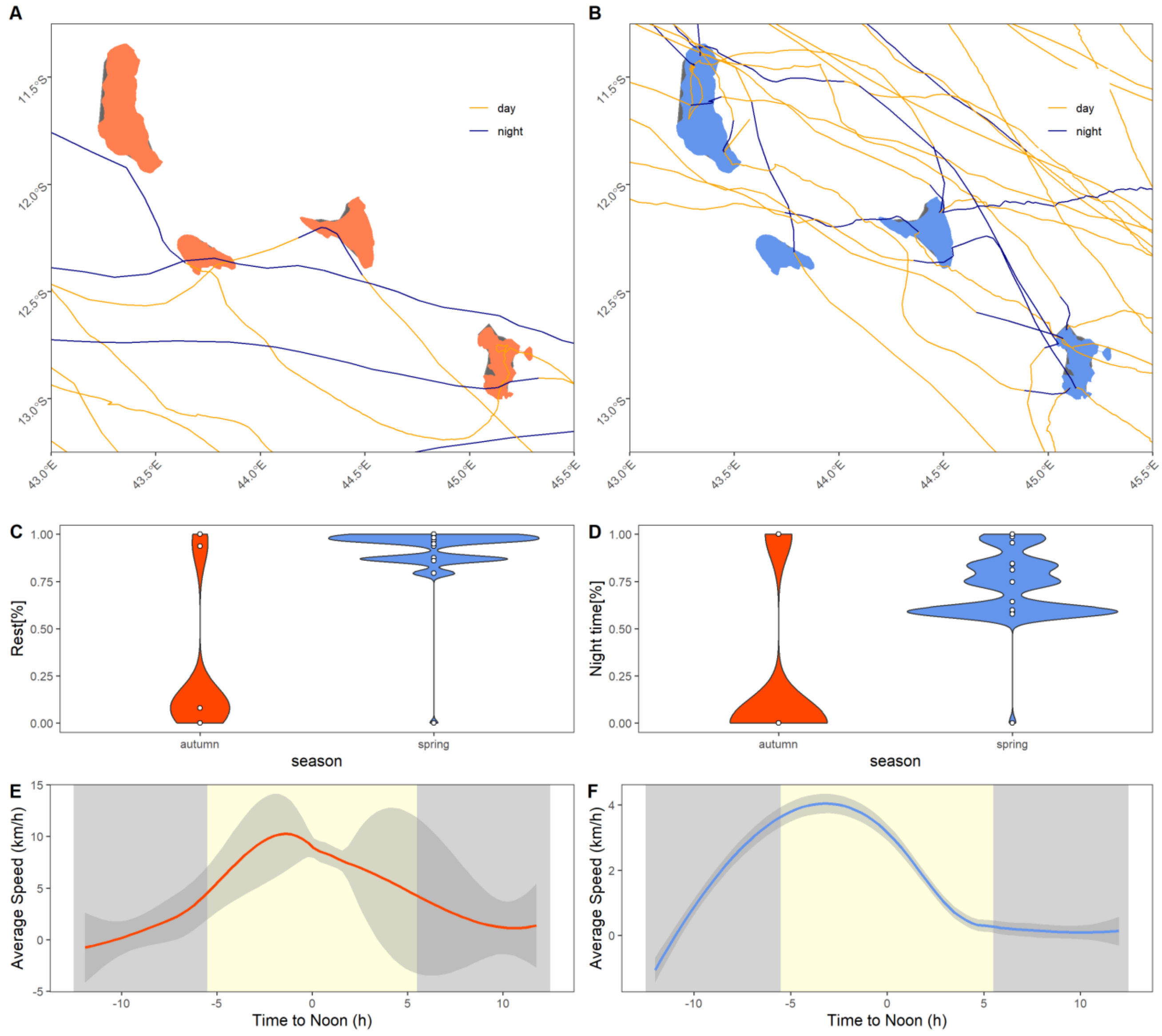


**Fig.S1.** Exploring activity patterns of Eleonora's falcons on islands. (A,B) Detailed maps showing tracks passing over or near the Comoros and Mayotte in autumn (A) and spring (B), respectively. (C,D) Violin plots showing (C) the proportion of time a falcon spent resting during an island visit and (D) the proportion of island visits that occurred at night. (E,F) Average ground speeds of falcons recorded over islands in relation to time of day. Background shading indicates the average daytime period (light yellow) and nighttime period (light gray).

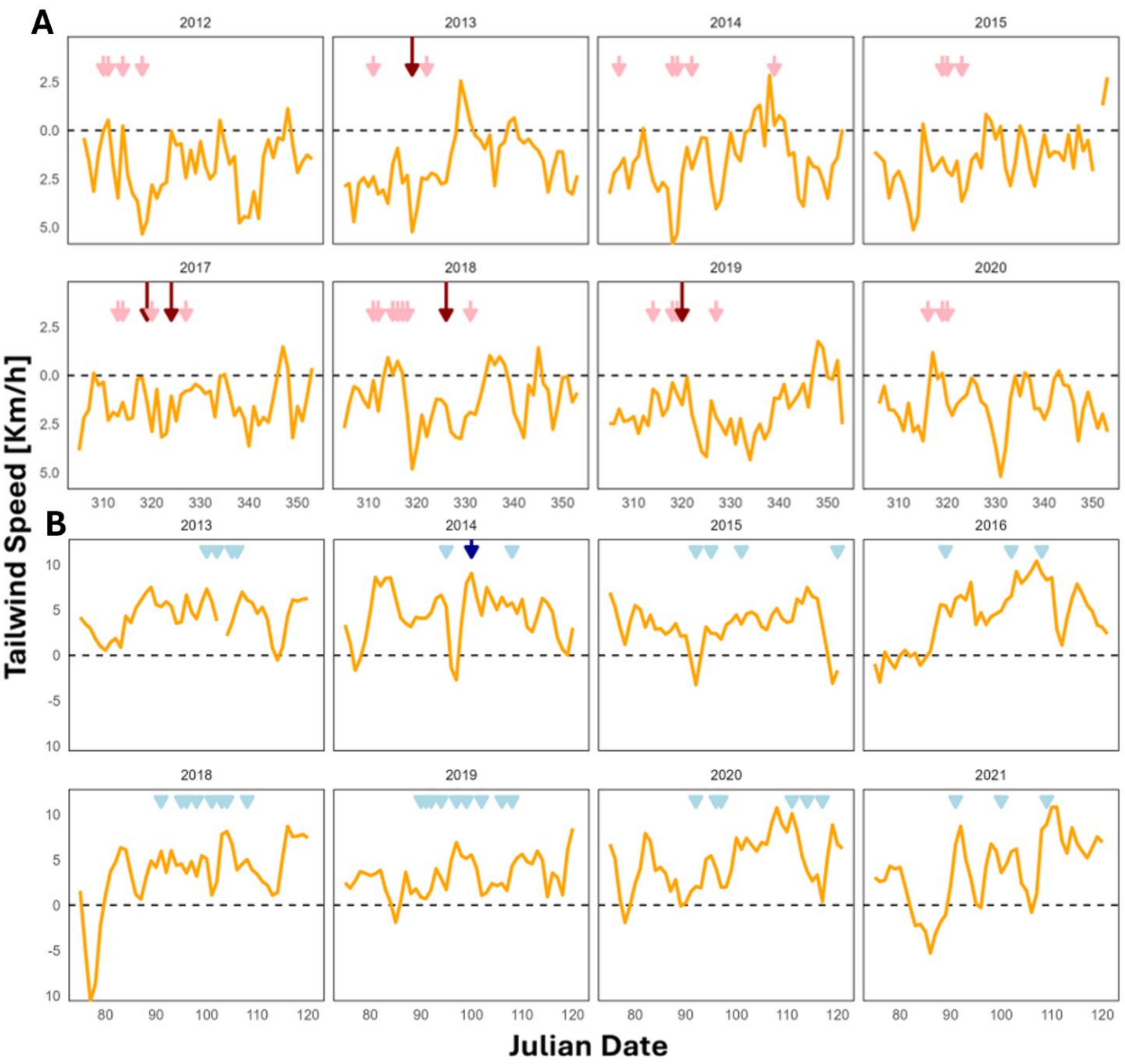


**Fig.S2** Dates of ocean-crossings by Eleonora's falcons in relation to daily tailwind conditions in the falcon's seasonal departure areas during (A) autumn and (B) spring, with each panel showing one of eight study years between 2012 to 2021. Orange lines indicate daily tailwinds across the falcons' seasonal departures areas, calculated relative to the falcons' average seasonal ocean-crossing direction. Coloured arrows indicate the dates on which crossings were recorded in autumn (red) and spring (blue). Lighter and shorter arrows indicate days on which one individual departed, whereas darker and longer arrows denote a higher number of departing individuals.
