## supplementarytables for "Falcons use wind assistance and remote islands to mitigate risk during ocean-crossings"

**Table S1.** Seasonal differences in ocean-crossing metrics (n = 83 crossings, 19 falcons), as estimated by generalized linear mixed models (GLMMs) with random intercepts per individual. Conditional $R^2$ could not be estimated for three response variables due to singular model fit, indicating that the inclusion of random intercepts resulted in over-fitting. In such cases we instead used standard linear models and only $R^2$ values are reported (models C, F, K). For the variable 'island count' (model F) we applied zero-inflated Poisson (ZIP).

| Figure | Response | $R^2$cond | $R^2$mar/ $R^2$ | Predictor | Estimate | Std. Error | df | t/z value | Pr(>\|t/z\|) |
|---|---|---|---|---|---|---|---|---|---|
| Fig.1C | Distance | NA | 0.81 | Autumn | 624.4 | 26.88 | 81 | 23.23 | **1.47E-37** |
| | | | | Spring | 707.94 | 38.25 | 81 | 18.51 | **7.88E-31** |
| Fig.1D | Duration | 0.7 | 0.69 | Autumn | 15.34 | 0.84 | 49.95 | 18.24 | **9.77E-24** |
| | | | | Spring | 15.68 | 1.15 | 69.14 | 13.68 | **2.42E-21** |
| Fig.1E | Duration Island | 0.21 | 0.13 | Autumn | 0.38 | 1.13 | 45.54 | 0.34 | 7.35E-01 |
| | | | | Spring | 5.38 | 1.44 | 68.31 | 3.73 | **3.93E-04** |
| Fig.1F | Islands Count | NA | 0.41 | Autumn | -1.45 | 0.52 | 79 | -2.76 | **5.86E-03** |
| | | | | Spring | 1.32 | 0.49 | 79 | 2.67 | **7.66E-03** |
| Fig.1G | Tail Wind speed | 0.43 | 0.42 | Autumn | -0.47 | 0.4 | 48.72 | -1.18 | 2.43E-01 |
| | | | | Spring | 4.29 | 0.55 | 68.21 | 7.77 | **5.53E-11** |
| Fig.1H | Ground Speed | 0.15 | 0.01 | Autumn | 42.29 | 1.09 | 42.7 | 38.98 | **5.15E-35** |
| | | | | Spring | 1.05 | 1.3 | 68.35 | 0.81 | 4.22E-01 |
| Fig.1I | Air Speed | 0.52 | 0.51 | Autumn | 47.38 | 1.24 | 50.26 | 38.26 | **7.80E-39** |
| | | | | Spring | -15.89 | 1.71 | 68.92 | -9.28 | **9.54E-14** |
| Fig.1J | Wind Displacement | 0.7 | 0.64 | Autumn | -91.49 | 30.13 | 37.67 | -3.04 | **4.32E-03** |
| | | | | Spring | 459.64 | 34.51 | 66.41 | 13.32 | **1.99E-20** |
| Fig.1K | Air Distance | NA | 0.23 | Autumn | 717.41 | 35.67 | 81 | 20.11 | **3.09E-33** |
| | | | | Spring | 247.83 | 50.76 | 81 | 4.88 | **5.17E-06** |

**Table S2**. Seasonal nocturnality differences in ocean-crossing metrics (n = 83 crossings, 19 falcons), as estimated by generalized linear mixed models (GLMMs) with random intercepts per individual.

| Figure | Response | $R^2$cond | $R^2$mar | Predictor | Estimate | Std. Error | df | t/z value | Pr(>\|t/z\|) |
|---|---|---|---|---|---|---|---|---|---|
| Fig.3D | Nocturnality [%h] | 0.06 | 0.04 | Autumn | 0.31 | 0.03 | 50.01 | 11.1 | **4.31E-15** |
| | | | | Spring | 0.08 | 0.04 | 68.75 | 1.94 | 5.69E-02 |
| Fig.3C | Nocturnal travel time [h] | 0.39 | 0.39 | Intercept | 5.12 | 0.7 | 55.61 | 7.26 | **1.32E-09** |
| | | | | Spring | 7.19 | 1 | 71.18 | 7.23 | **4.47E-10** |

**Table S3.** Results of generalized linear models (GLMs) testing the effects of wind speed during ocean crossings and the start latitude of ocean-crossings on the number of islands used by Eleonora's falcons during autumn and spring. Models were fitted using a Poisson error distribution with a log-link. We used quasi-AIC (QAIC) to identify the best model for each season (bolded QAIC). Pseudo-$R^2$ (McFadden's) are provided to assess explanatory power of the models.

| response | Season | Estimate | Std. Error | z value | Pr(>\|z\|) | Predictor | $R^2$ | QAIC |
|---|---|---|---|---|---|---|---|---|
| Island use | Spring | -0.18 | 0.34 | -0.52 | 6.03E-01 | Intercept | 0.07 | 75.49 |
| | | -0.10 | 0.07 | -1.35 | 1.77E-01 | Tailwind | | |
| | | -7.95 | 2.77 | -2.87 | **4.09E-03** | Intercept | 0.24 | 76.31 |
| | | -0.54 | 0.20 | -2.73 | **6.37E-03** | Start latitude | | |
| | | 0.89 | 4.80 | 0.19 | 8.52E-01 | Intercept | 0.39 | **58.22** |
| | | 0.09 | 0.36 | 0.27 | 7.90E-01 | Start latitude | | |
| | | -2.48 | 1.20 | -2.07 | **3.84E-02** | Tailwind | | |
| | | -0.18 | 0.09 | -2.02 | **4.34E-02** | Start latitude*Tailwind | | |
| | | -7.14 | 2.82 | -2.53 | **1.14E-02** | Intercept | 0.27 | 76.08 |
| | | -0.51 | 0.20 | -2.55 | **1.09E-02** | Start latitude | | |
| | | -0.08 | 0.08 | -1.00 | 3.16E-01 | Tailwind | | |
| | Autumn | -1.68 | 0.44 | -3.82 | **1.34E-04** | Intercept | 0.06 | **30.44** |
| | | 0.19 | 0.19 | 1.04 | 2.99E-01 | Tailwind | | |
| | | 21.23 | 7.90 | 2.69 | 7.18E-03 | Intercept | 0.74 | 71.84 |
| | | 1.83 | 0.67 | 2.74 | 6.22E-03 | Start latitude | | |
| | | 20.02 | 7.18 | 2.79 | 5.29E-03 | Intercept | 0.78 | 73.93 |
| | | 1.70 | 0.61 | 2.77 | 5.54E-03 | Start latitude | | |
| | | 1.49 | 3.35 | 0.44 | 6.57E-01 | Tailwind | | |
| | | 0.10 | 0.27 | 0.38 | 7.03E-01 | Start latitude*Tailwind | | |
| | | 19.38 | 6.71 | 2.89 | 3.87E-03 | Intercept | 0.77 | 72.30 |
| | | 1.65 | 0.57 | 2.89 | **3.88E-03** | Start latitude | | |
| | | 0.22 | 0.22 | 0.99 | 3.24E-01 | Tailwind | | |

**Table S4.** Results of the GLM examining whether the proportion of resting time on islands was correlated with the amount of time spent on an island at night, by day, or the total duration of island visits. We assumed a beta error distribution using a logit link function. Each model includes only one predictor. For clarity, intercepts are reported only for the null models. $R^2$ values reflect the variance explained by the fixed effects.

| Response | Season | Predictor | Estimate | Std. Error | z value | Pr(>\|z\|) | $R^2$ | AIC |
|---|---|---|---|---|---|---|---|---|
| rest [%] | spring | Intercept (null model) | 1.88 | 0.03 | 64.87 | 0.00E+00 | 0.00 | -6872.04 |
| | | Intercept | -0.82 | 0.07 | -11.29 | **1.55E-29** | 0.54 | **-7476.03** |
| | | nighttime duration | 0.29 | 0.01 | 35.81 | **7.22E-281** | | |
| | | Intercept | 2.14 | 0.04 | 50.68 | 0.00E+00 | 0.12 | -6943.05 |
| | | daytime duration | -0.06 | 0.01 | -8.59 | **8.74E-18** | | |
| | | Intercept | 2.33 | 0.12 | 19.23 | 2.09E-82 | 0.05 | -6884.88 |
| | | total duration | -0.03 | 0.01 | -3.91 | **9.36E-05** | | |
| | autumn | Intercept (null model) | -0.44 | 0.09 | -4.98 | 6.35E-07 | 0.00 | -176.00 |
| | | Intercept | -2.10 | 0.08 | -27.76 | 1.28E-169 | 0.98 | -406.58 |
| | | nighttime duration | 1.01 | 0.06 | 17.67 | **6.83E-70** | | |
| | | Intercept | 2.48 | 0.14 | 17.96 | 4.20E-72 | 0.93 | **-543.43** |
| | | daytime duration | -1.87 | 0.07 | -26.20 | **2.66E-151** | | |
| | | Intercept | -1.95 | 0.15 | -13.03 | 8.35E-39 | 0.83 | -300.54 |
| | | total duration | 0.38 | 0.04 | 10.49 | **9.75E-26** | | |
